# Development and characterization of a human model of arteriovenous malformation (AVM)-on-a-chip

**DOI:** 10.1101/2022.01.20.477166

**Authors:** Kayla Soon, Mengyuan Li, Ruilin Wu, Williamson D. Turner, Joshua D. Wythe, Jason E. Fish, Sara S Nunes

## Abstract

Brain arteriovenous malformations (AVMs) are a disorder wherein abnormal, enlarged blood vessels connect arteries directly to veins, without an intervening capillary bed. AVMs are one of the leading causes of hemorrhagic stroke in children and young adults. Most human sporadic brain AVMs are associated with genetic activating mutations in the *KRAS* gene. Our goal was to develop an *in vitro* model that would allow for simultaneous morphological and functional phenotypic data capture in real time during AVM disease progression. By generating human endothelial cells harboring a clinically relevant mutation found in most human patients (activating mutations within the small GTPase *KRAS*) and seeding them in a dynamic microfluidic cell culture system that enables vessel formation and perfusion, we demonstrate that vessels formed by *KRAS4A^G12V^* mutant endothelial cells (ECs) were significantly wider and more leaky than vascular beds formed by wild-type ECs, recapitulating key structural and functional hallmarks of human AVM pathogenesis. Immunofluorescence staining revealed a breakdown of adherens junctions in mutant KRAS vessels only, leading to increased vascular permeability, a hallmark of hemorrhagic stroke. Finally, pharmacological blockade of MEK kinase activity, but not PI3K inhibition, improved endothelial barrier function (decreased permeability) without affecting vessel diameter. Collectively, our studies describe the creation of human KRAS-dependent AVM-like vessels *in vitro* in a self-assembling microvessel platform that is amenable to phenotypic observation and drug delivery.

## Introduction

A brain arteriovenous malformation (AVM) is an abnormal tangle of blood vessels in the brain, wherein feeding arteries are directly connected to draining veins, without an intervening capillary bed, characterized by high blood flow, and dilated, tortuous vessels ^1^. Instead of a normal hierarchical vascular network, this abnormal vascular conduit, or shunt, lacks the integrity and stability characteristic of normal capillary networks. Individuals with AVMs are at increased risk of hemorrhage as patients age due to the direct shunting of high-pressure flow from the artery ^2,3^. AVMs are responsible for 4% of all intracerebral haemorrhages and account for almost one third of brain bleeds in young adults ^4^. Currently, there are no approved pharmacological treatments for brain AVMs, and the only available treatment options for patients are surgical resection and embolization of small AVMs or stereotactic radiosurgery for inoperable vessels to reduce vessel size. Both procedures are highly invasive with a recurrent hemorrhage risk as high as 18% in the first year. Approximately one third of all AVM cases are untreatable with currently available options ^5^.

Various mutations within genes encoding proteins in the mitogen-activated protein kinase (MAPK/ERK, e.g. the RAS-RAF-MEK-ERK) pathway have been implicated in AVM pathology. Recently, somatic activating KRAS mutations in endothelial cells (ECs) have been reported in as many as 50% of clinical samples ^6,7^. Nikolaev *et al*. ^7^ showed that somatic activating KRAS mutations in ECs derived from patient AVMs, or expression of mutant KRAS in human umbilical vein endothelial cells (HUVECs), leads to increased ERK1/2 phosphorylation and disruption of vascular endothelial (VE)-cadherin localization in adherens junctions. This is thought to increase the permeability of the vessel wall, and to leave vessels prone to hemorrhage ^7,8^. Although these genetic mutations are sufficient for the development of AVM in zebrafish and mouse models ^9^, the initiation, development, and remodelling of these dynamic vessel malformations are influenced by factors in the microenvironment. Specifically, vascular endothelial growth factor (VEGF) and other pro-angiogenic factors are highly expressed in AVMs, leading to increased vessel permeability and susceptibility to rupture ^10,11^. Other pathways, such as Notch and transforming growth factor (TGF)-β/bone morphogenetic protein (BMP) have also been implicated in AVM regulation ^12^, particularly with discontinuity of cell-cell junctions and cell proliferation, resulting in vascular dysplasia and instability ^11,13,14^. Moreover, growing evidence suggests that the AVM microenvironment may promote AVM growth. Inflammation driven by local macrophages and reduction of pericyte coverage have been described as contributing factors in AVM pathogenesis, as they promote excessive angiogenic responses ^15,16^. Additionally, a high flow rate is a defining feature in AVMs that leads to a positive feedback loop to increase local vessel enlargement in high-velocity vessels ^17^. However, there is limited knowledge as to how environmental cues from surrounding cells, hemodynamics, and the KRAS mutation contribute to the pathogenesis of AVMs.

Animal models of the disease have achieved variable degrees of success in generating vessel lesions with characteristics that match human pathology. Several transgenic mouse models of hereditary hemorrhagic telangiectasia (HHT), an inherited disease known to cause AVMs, involve genetic inhibition or deletion of either *Eng* or *Alk1* genes in combination with angiogenic induction to generate AVMs ^18–20^. While early postnatal, endothelial specific deletion of either ALK or ENG in mice produces brain AVMs, deletion in adults fails to produce brain AVMs in the absence of angiogenic stimuli ^21,22^. Critically, successful murine models have recapitulated key features of AVM, such as vessel dilation, arteriovenous shunts, and hemorrhaging ^23^. The incidence of observable AVMs can be low, and there is often high and early mortality associated with these models ^23,24^. Recently, mouse and zebrafish models with the KRAS^G12V^ or KRAS^G12D^ mutation displayed enlarged, tortuous vessels that may be prone to hemorrhage ^9,25^. These animal models have contributed to defining a mechanism that may underlay the majority of sporadic brain AVMs, as these activating mutations are detected in around 50% of sporadic bAVMs. However, these models are costly, in terms of both time and resources, and have yet to be validated in a human-relevant model other than monolayer cell culture studies ^7,9,26^.

In recent years, organ-on-a-chip models have emerged as candidates to address the limitations of 2D models and species-specific factors in disease modeling and drug development. In comparison to cell monolayers, organ-on-a-chip models have increased architectural complexity, mimicking key aspects of *in vivo* tissue physiology ^27^. Using a combination of adjoining channels and interfaces, they enable precise spatiotemporal control of mechanical and chemical stimuli to probe tissue responses ^28,29^. Several studies have successfully produced vessel-like networks using ECs from different origins, in combination with supporting cells such as fibroblasts ^30,31^, pericytes ^32,33^, or mesenchymal stem cells ^34^. Self-assembled microvessels on-chip models are formed in the presence of media flow, allowing vessel perfusion and mimicking in vivo flow dynamics and feedback ^35^. Rather than having predefined geometries, the self-assembling nature of microvasculature-on-a-chip forms an organic vasculature with vessels of varying shapes and sizes. This aspect is particularly relevant when studying diseases that lead to alterations in vessel diameter, such as in AVMs ^29^.

Our goal is to recapitulate the key hallmarks of AVMs *in vitro* using human ECs harbouring a clinically relevant KRAS mutation described in patients with sporadic AVMs in a microfluidic platform that allows 3D vessel assembly and perfusion. We also aim to test the impact of MEK inhibitors on AVM structural and functional parameters. We found that a mosaic culture of KRAS^G12V^ mutant and wildtype ECs led to irregularly shaped and enlarged microvessels, with increased vessel wall permeability. Treatment with a MEK inhibitor restored barrier function but did not reverse vessel dilation.

## Materials and Methods

### 2.1 Cell Culture

Immortalized (Im)-HUVECs (a gift from Dr. Roger Kamm at MIT) were grown in VascuLife VEGF Endothelial Medium (Lifeline Cell Technology). Im-HUVECs constitutively express green fluorescent protein (GFP) ^36^. Primary HUVECs and KRAS^G12V^ Im-HUVECs were grown in Endothelial Cell Medium (ScienCell). Normal human lung fibroblasts (NHLF; Lonza) were grown in DMEM (ThermoFisher) supplemented with 20% fetal bovine serum (FBS). NHLF of passage 5-7 were used in experiments. All cells were cultured in cell culture flasks coated with 1% gelatin until they reached 85-90% confluency.

### 2.2 Generation of Stable, Tet-On KRAS4A^G12V^ HUVECs

Telomerase-immortalized human umbilical vein endothelial cells (Im-HUVECs, or hTERT-HUVECs) were kindly provided by Dr. Arnold Hayer (McGill University) ^37^. A piggyBac ITR flanked plasmid, pB-TA-ERN - a gift from Knut Woltjen (Addgene plasmid # 80474) ^38^ was combined with pDONR221-V5-mScarlet-I-hsKRAS4A^G12V^ (addgene #156406) ^9^ and LR recombinase to generate pB-TetOn-mScarlet-KRAS4A^G12V^ by Gateway recombination. To generate the stable Im-HUVEC tunable KRAS^G12V^ line, 1.5 μg of pB-TetON-mScarlet-KRAS4A-G12V (JDW 891) and 1 μg of a plasmid encoding a hyperactive *piggyBac* transposase (hyPBase) ^39^ were co-electroporated into Im-HUVEC using the P5 Primary Cell Kit (Lonza) and a 4D-nucleofector (Lonza). The program used was CA167. After electroporation, 100 μL of FBS was immediately added to the cuvette and incubated at 37°C for 5 minutes. Cells were transferred to a 12-well plate after incubation and grown for 2 days in antibiotic-free media. For antibiotic selection, 0.3 mg/mL G418 was added for 5 days with media change every other day. After selection, cells (KRAS4A^G12V^ HUVECs) were expanded and passaged.

### 2.3 Device Fabrication

AutoCAD was used for creating the device design (supplemental Figure 1A), which was written onto a chrome photomask (Heidelberg upg501 Mask Writer). The standard lithography process supplied by the photoresist manufacturer was used to generate the silicone molds. Briefly, SU-8 2050 photoresist (MicroChem) was spun onto a silicon mold to achieve a 100 μm thick layer. The photomask was used to polymerize (240 mJ/cm^2^ exposure energy) the pattern onto the wafer. An optical profiler was used to confirm surface and edge finishes of each silicon wafer mold. Silicone mold heights ranged from 90-120 μm.

PDMS was mixed at a 1:10 ratio (elastomer to base) and left to de-gas for 20 min. PDMS was poured onto the wafers inside an acrylic ring so that the final PDMS mold was easily removable. PDMS was baked for 3 hours in the oven at 80°C. After baking, the PDMS was left overnight to cool before removing the PDMS mold. This was to prevent the SU-8 features from peeling off while removing the PDMS. Access holes of 2 mm for the side channels and 1 mm for the tissue chamber were cut out using biopsy punches. PDMS coated glass slides were prepared by spreading 2 mL of PDMS onto a 75 x 50 mm glass slide and baking for 1 hr. Prior to bonding, PDMS molds and PDMS-coated glass slides were covered with packing tape to prevent dust accumulation. The devices were bonded together using plasma treatment (Harrick Plasma). Media reservoirs were created by attaching cryovials that had their bottoms cut out using PDMS (supplemental Figure 1B). The entire device unit was autoclaved before use (supplemental Figure 1C).

### 2.4 Cell Seeding

Twenty-four hours after KRAS4A^G12V^ HUVECs were plated into cell culture flasks, doxycycline (Sigma-Aldrich, D3072) was added to the media (final concentration: 5 μg/mL) to induce KRAS4A^G12V^ expression (Supplemental Figure 2). Im-HUVECs and NHLF were seeded in fibrin gels (initial concentration: 10 mg/mL bovine fibrinogen) at a 4:1 ratio (Im-HUVECs to NHLF) for a concentration of 10 million total cells/mL. To generate AVM-like microvessels, KRAS^G12V^ Im-HUVECs were combined with a Im-HUVEC:NHLF cell suspension at the described ratios. The cell suspension was gently pipetted into one side of the tissue chamber until the chamber was filled. Devices were placed in the incubator for 15 min for gel polymerization, after which Im-HUVECs were seeded to the side channels coated with 0.1% gelatin in PBS (BioShop cat# GEL771.500) for 2 hrs in Vasculife media. During this time, devices were routinely checked, and bubbles removed. To establish perfusion, Vasculife media was added in the following volumes to the media reservoir #1: 1500 μL, #2: 1000 μL, #3: 500 μL, #4: 200 μL to achieve an approximate pressure gradient of #1: 23 mm H_2_O, #2: 18 mm H_2_O #3: 8 mm H_2_O, #4: 3 mm H_2_O, as previously reported ^31^. Media reservoirs were refilled every 24 hours. The media flow volume across the tissue chamber was reversed (1&3 ↔ 2&4) every 2 days to prevent ECs from migrating to side channels (Figure 1A).

**Figure 1:**
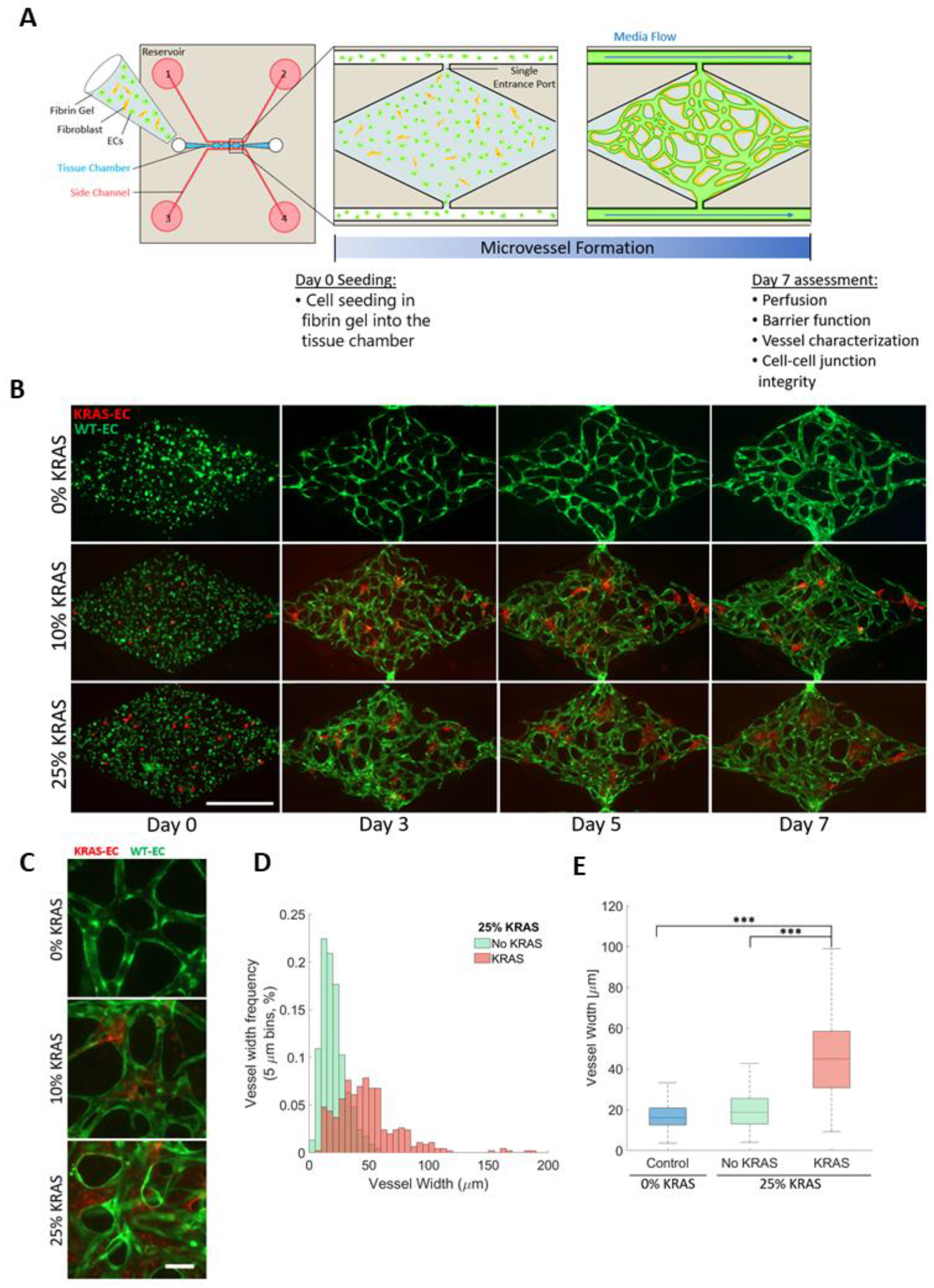
Generation of a perfused human AVM-like vasculature-on-a-chip. (A) Schematic representation of the experiments and timelines. (B) Representative time course of vessel network formation using different ratios of wild-type (WT)- and KRAS4A^G12V^-ECs. Images of cell suspension immediately after seeding on day 0. WT-ECs (green) and KRAS4A^G12V^ ECs (red) are round and evenly distributed within the fibrin gel. By day 3, tube-like structures begin to form as they migrate towards each other. By day 5, microvessel connections and network shape begins to stabilize. By day 7, the microvessel network is fully interconnected, lumenized and perfused. (Scale bar 500 μm). (C) Zoomed fluorescent images of microvessel segments of 0% KRAS4A^G12V^ (control), 10% KRAS4A^G12V^, 25% KRAS4A^G12V^ vasculatures (scale bar, 50 μm) (D) Distribution of microvessel width measurements in 25% KRAS4A^G12V^ vasculatures between segments without KRAS4A^G12V^ ECs (green) and segments with some visible KRAS4A^G12V^ (red) (bin size: 10 μm). KRAS4A^G12V^ positive vessel distribution is much wider and variable compared to vessel with no KRAS4A^G12V^ (E) Quantification of the average vessel diameter in microvessel networks in 0% KRAS4A^G12V^ ECs (blue) compared to 25% KRAS4A^G12V^ microvasculatures with (red) and without (green) KRAS4A^G12V^ ECs. (One-way ANOVA with Tukey’s multiple comparison test; ***p < 0.001, N=3; mean ± SD)

### 2.5 Microvessel Network Perfusion and Permeability

To assess microvessel perfusion and permeability, 1 mg/mL of 70 kDa dextran conjugated to rhodamine b isothiocyanate (Rhod B, Sigma) was introduced from one media reservoir. Once the dextran solution had perfused the entire vessel network (t=1) images of each diamond within the tissue chamber were acquired at 10 min intervals, for 30 min total. Images were acquired using a Zeiss AxioObserver Widefield microscope within a 37°C temperature controlled incubation chamber (5% CO_2_).

### 2.7 Immunofluorescence Staining

Immunostaining was performed *in situ*, by perfusing all reagents into the device through the side channels. The devices were fixed using 4% paraformaldehyde (PFA) for 20 min at room temperature and washed with PBS overnight. Devices were then permeabilized with 0.05% Tween 20 in PBS and washed for 2-3 hrs with PBS. Then, a blocking solution (5% BSA) was added and incubated overnight at 4°C. VE-cadherin primary antibody (Santa Cruz cat# sc-9989, 1:200 in blocking solution) was incubated overnight at 4°C. Devices were washed with PBS for 4-6 hrs and subsequently incubated with Alexa Fluor 647-conjugated goat anti-mouse IgG (H+L) (ThermoFisher cat# A32728, 1:300) and Hoechst 33342 (Sigma-Aldrich cat# B2261, 1:10000) in blocking solution overnight at 4°C. Devices were washed with PBS and imaged using the Zeiss AxioObserver Widefield microscope.

For immunofluorescence staining of cell monolayers, primary HUVEC and Im-HUVEC cells were washed with PBS and fixed with 4% paraformaldehyde at room temperature for 20 minutes. Cell were then permeabilized with 0.5% Triton X-100 for 10 minutes, followed by blocking with 5% BSA for 1 hour at room temperature. Primary antibodies were added to the cells and incubated at 4°C overnight. Primary antibodies used were VE-cadherin (Santa Cruz cat# sc-9989; Mouse; 1:200 dilution) and Phospho-p44/42 MAPK (Erk1/2) (Thr202/Tyr204) (Cell Signaling Technology cat# 9101S; Rabbit; 1:200 dilution). On the following day, primary antibodies were removed and cells were washed with PBST (0.1% Triton X-100 in PBS). Secondary antibodies were incubated at room temperature for 2 hours, covered in foil. Secondary antibodies used were Alexa 488 goat anti-mouse (Invitrogen cat# A11001; 1:200 dilution) and Alexa 647 goat anti-rabbit (Invitrogen A21245; 1:200 dilution). After secondary antibody incubation, cells were washed with PBST and Hoechst (ThermoFisher cat# 62249) staining was done at 1:500 dilution for 10 minutes at room temperature. Then, cells were washed with PBS and mounted with ProLong^™^ Gold Antifade Mountant (Invitrogen cat# P36930). Confocal imaging of primary HUVEC and Im-HUVEC cells grown in 2D was done with an Olympus Fluoview 1000 Confocal microscope Olympus IX81 inverted stand (Olympus, CA, USA) using the Plan Apo 40x/1.35 NA oil immersion objective (NA 1.3). Excitation wavelengths were 405nm for Hoechst, 473nm for Alexa Fluor 488, 559nm for mScarlet and 635nm for Alexa Fluor 647. Image processing was done uniformly across all conditions with the FV10-ASW 4.2 Viewer (Olympus, CA, USA) and FIJI (v2.1.0/1.53c).

### 2.8 Treatment of Microvessel Network

For drug treatment experiments, 25% KRAS^G12V^ vasculatures were treated with 1 μM MEK inhibitor (MEKi, SL327, Sigma) or 10 μM PI3K inhibitor (PI3Ki, LY294002, Sigma) in Vasculife media on days 3 or 5. Treatment was maintained until endpoint (day 7). DMSO was used as a vehicle control.

### 2.9 Image Analysis

All images were analysed using Zen blue (ZEISS) and ImageJ (FIJI). All graphs and calculations were done using MATLAB (MathWorks, R2020). Angiotool ^40^ was used to assess total branch length. Diameter, junction, and branch measurements were done manually using Zen blue. To measure fluorescent intensity in perfusion assays, a binary mask was created from the FITC channel. Two separate images were generated for the intravascular and extravascular perfusion by using the AND operation of the mask and inverse mask over the TRITC channel. The intensity values were then averaged from the intravascular and extravascular images respectively. The permeability coefficient (P) was calculated using the previously described equation ^41^.

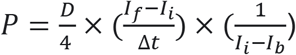

Where D is the average diameter, Δ*t* is the time interval, *I_f_*, *I_i_*, *I_b_* is the fluorescence intensity of the final intravascular, initial intravascular, and background (extravascular) intensity respectively.

### 2.10 Statistical Analysis

All statistical analyses were performed using Prism 9 software (GraphPad). All data presented are mean ± standard deviation (SD). One-way ANOVA with Tukey’s multiple comparisons was used to assess significance between 3 groups. The paired two-tailed t-test was used to assess diameter changes over time. The unpaired two-tailed t-test was used to assess significance between 2 groups. A p value <0.05 was considered significant. Each diamond within the tissue chamber was considered a technical replicate (indicated by n). Separate experiments (indicated by N) were considered biological replicates. At least 3 ‘N’ were used for each experimental/seeding condition.

## Results

### 3.1 Generation of perfused human AVM-like vasculatures *in vitro*

We used a previously described microfluidic device consisting of a central tissue chamber divided into three diamond-shaped chambers, with one port on each end connected to side channels where media is delivered ^31,35^ (Figure 1A). The 50 μm ports create a single entry and exit in the 1×2 mm diamond chamber and segments the vasculature into three technical replicates (Figure 1A and Supplemental Figure 1A). In this platform, perfusable microvessels were reliably generated *in vitro* over the course of 7 days through self-assembly of endothelial cells (i.e., Im-HUVECs), supported by fibroblasts at a 4:1 ratio. Additionally, the inclusion of endothelial cells in the side channels, generating a monolayer, helped with vessel anastomosis and prevented leakage ^35^ (Figure 1A). To prevent cells from migrating out of the tissue chamber, the media flow direction was switched between the two side channels every two days by switching inlet function between reservoirs 1&3 to 2&4 (Figure 1A).

For cell seeding (day 0), cells were loaded into the tissue chamber in a fibrin gel. By day 3, ECs extended to neighbouring cells and clustered together to form rudimentary vessels. By day 5, the network structure was clearly established and interconnected. At this time point, microvessels were marginally perfusable (not shown), indicating that the network was lumenized. By day 7, microvessels expanded in diameter to create interconnected, well-perfused, and stable vessel networks (Figure 1B, top panel).

To create the AVM disease model, ECs containing a stably integrated, doxycycline-inducible mScarlet (mScar)-KRAS4A^G12V^ expression construct were induced prior to seeding into the microfluidic device ^9^. Expression was confirmed by the presence of mScar after the addition of doxycycline (Supplemental Figure 2). Induction of KRAS4A^G12V^ elevated pERK levels in the mutant ECs, indicating functional KRAS protein activity and enhanced MAPK signaling. Consistent with our previous studies ^5,7^, mutant KRAS expression led to a down-regulation of VE-cadherin at cell junctions (Supplemental Figure 2). KRAS4A^G12V^ ECs induced with doxycycline were combined with WT-ECs, at 10% or 25% of the total EC number, in fibrin gels and seeded into the microfluidic devices.

WT-ECs seeded without KRAS4A^G12V^ ECs served as controls. Over the course of 7 days, heterogeneous KRAS4A^G12V^ and WT-EC co-cultures effectively self-assembled into perfusable microvessels (Figure 1B, middle and bottom panels). In devices containing KRAS4A^G12V^ ECs, mScar-positive segments were present within the vessel network, confirming that the KRAS4A^G12V^ ECs had integrated into the vasculature (Figure 1B, C). As expected, vasculatures with KRAS4A^G12V^ ECs had distinct aberrant vessel structures. Vascular networks containing 25% KRAS4A^G12V^ ECs resembled vascular plexuses rather than the typical microvascular networks formed using WT-ECs (Figure 1B, C).

To assess potential changes in vessel diameter associated with mutant KRAS expression, the width of individual microvessels in the 25% KRAS vasculatures were measured on day 7. Vessel width in the 25% KRAS microvessels was larger and highly heterogenous compared to 0% KRAS, resulting in a larger distribution (Figure 1D). In line with human brain AVMs, we found that vessel segments in 25% KRAS microvessels with KRAS4A^G12V^ ECs were significantly wider than vessel segments within the same network that lacked KRAS4A^G12V^ ECs, measuring 47.3 ± 28.5 μm and 18.2 ± 7.2 μm, respectively (Figure 1E). There was no significant difference between vessel widths in mixed cultures in sections without KRAS and the vessel width in wild-type monocultures (i.e., 0% KRAS) (17.5 ± 7.3 μm) (Figure 1E). This demonstrated that vessel enlargement only occurs where KRAS4A^G12V^ ECs are present, but this does not affect surrounding vessels formed by WT-ECs within the same network. Unfortunately, it was not possible to assess vessel width of KRAS4A^G12V^ EC vessels in the 10% KRAS networks, as KRAS4A^G12V^ ECs mainly concentrated in vessel junctions when seeded at low ratios (Figure 1C, center image). Hereafter, vasculatures generated with 0%, 10%, and 25% KRAS4A^G12V^ ECs at seeding will be referred to as 0%, 10% and 25% KRAS vasculatures respectively.

### 3.2 Changes in vessel network parameters towards AVM-like morphology

To assess the morphological and functional changes in vessel networks formed with KRAS4A^G12V^ ECs, we evaluated branch length, junction number, and vessel area coverage. As a result of vessel enlargement, the total area covered by the vessel network is greater in 10% and 25% KRAS microvessels (50.0 ± 3.2% and 53.4 ± 3.5%, respectively), and significantly higher than 0% KRAS microvessels (42.1 ± 6.9%) (Figure 2A).

**Figure 2:**
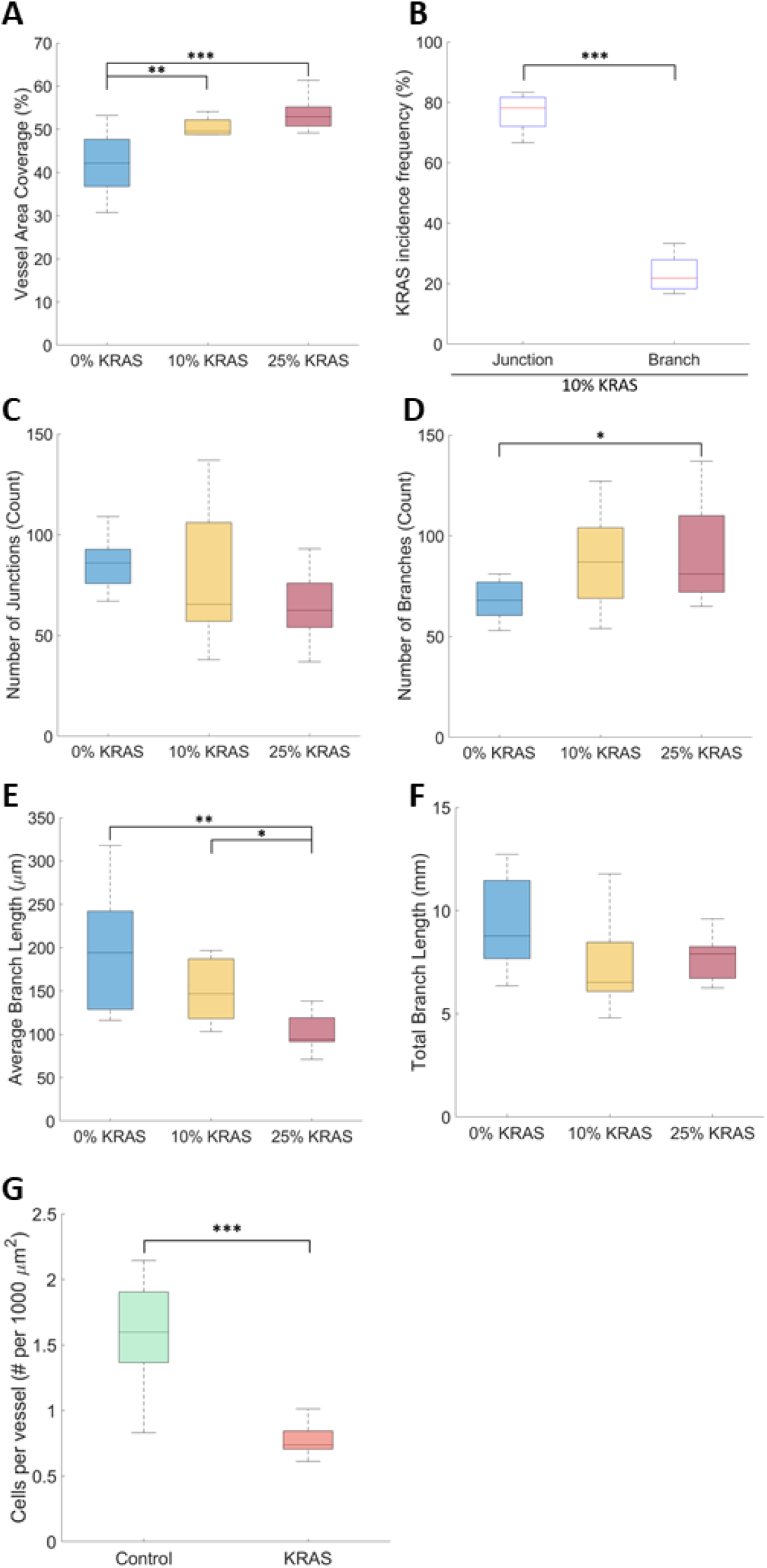
Quantification of AVM-like microvessel morphological characteristics. Quantification of (A) area coverage of 0% (blue), 10% (yellow), and 25% KRAS4A^G12V^ (red) microvasculatures. (One-way ANOVA with Tukey’s multiple comparison test; **p < 0.01, ***p < 0.001, N=3; mean ± SD) (B) Incidence rate of KRAS4A^G12V^ located in vessel junctions (left) compared to branches (right) in 10% KRAS4A^G12V^ microvasculatures. (Unpaired Student’s t-test; ***p < 0.001, N=3; mean ± SD). Quantification of (C) number of junctions, (D) number of branches, (E) average branch length, and (F) total branch length of 0% (blue), 10% (yellow), and 25% KRAS4A^G12V^ (red) microvasculatures. (One-way ANOVA with Tukey’s multiple comparison test; *p < 0.05, **p < 0.01, N=3; mean ± SD) (G) The number of cells counted in GFP^+^ (green) or mScarlet^+^ (red) vessel segments normalized to 1000 um^2^ vessel area. (Unpaired Student’s t-test; ***p < 0.001, N=3; mean ± SD)

We next visualized where the mutant KRAS cells localize in the network. In 10% KRAS vasculatures, KRAS4A^G12V^ ECs preferentially localized in vessel junctions at a significantly higher rate than expected if this was a random process, and were depleted at branches, as shown in Fig. 1C and quantified in Fig. 2B. Seventy eight ± 9.0% of KRAS4A^G12V^ ECs localized at junctions, compared to 22.0±9.0% localizing at vessel branch segments. Despite the increased frequency of mutant KRAS-ECs at the junctions, the number of junctions in each vasculature was not significantly different (Figure 2C). Conversely, there was a significant increase in the number of branches between 0% KRAS (67.5±9.7) and 25% KRAS (90.0±24.2) vasculatures (Figure 2D), indicating that there were more branches extending from a single junction.

The average branch length in the overall network decreased from 190.1±70.0 μm in 0% KRAS to 160.7±58.3 μm in 10% KRAS and 101.8±20.1 μm in 25% KRAS vasculatures (Figure 2E). As a result of the inverse relationship between average branch length and count, the total branch length, being the length of all the vessel segments end-to-end, was not significantly different. The total branch length was 9.4±2.3, 7.2±2.0, and 7.8±1.1 mm in the 0%, 10%, and 25% KRAS microvessels, respectively (Figure 2F). Taken together, these measurements indicate that the inclusion of KRAS4A^G12V^ ECs in microvessel formation affected the overall vessel networks by shortening branch length and increasing vessel density.

By counting the number of cells in vessel segments with only GFP or mScar cells, we determined that the average cell density in vessels made from WT-ECs was two times higher than that of mScar KRAS4A^G12V^-ECs (1.7 and 0.8 cells/1000 μm^2^, respectively). This suggests that the resulting enlargement of microvessels in 10% and 25% vasculatures is not the result of an excessive number of ECs, but rather it is due to an increase in the overall size of KRAS4A^G12V^ ECs (e.g. hypertrophy, rather than hyperplasia) (Figure 2G).

### 3.3 AVM-like vessels display increased permeability

To test whether KRAS microvessels exhibited weakened barrier integrity, we assessed the extravasation of 70 kDa dextran conjugated to rhodamine b isothiocyanate (rhod B). On day 7, dextran-rhod B was added to the inlet and allowed to perfuse throughout the vasculature (denoted as time = 1 min). Images collected every 10 mins for a time course of 30 mins after initial dextran perfusion, confirmed that the control microvessel network was intact and lumenized, showing physiological barrier properties. Tthis also demonstrated that vessels were well connected to the inlets and outlets of the tissue chamber. No focal leakage was observed (Figure 3A, top panel). In addition, we observed an increase in interstitial fluorescence intensity over time in both 10% and 25% KRAS vasculatures, demonstrating increased permeability (Figure 3A, middle and bottom panel). In 10% KRAS microvessels, rhod B dextran was observed leaking from a singular point, originating from the vessel junction in 15% of the devices (Figure 3A, middle panel). In the other 10% KRAS vasculatures there was no apparent source of leakage. Instead, an overall increase in interstitial fluorescence intensity was observed. In the 25% KRAS microvessels, the overall vasculature was more permeable to rhod B dextran and there was progressive leakage throughout the chamber due to the widespread distribution of KRAS4A^G12V^ ECs throughout the vascular network (Figure 3A, bottom panel). No focal leakage points were observed. Quantification of the fluorescence intensity demonstrated a significant increase in extravascular Dextran-rhod B in 25% KRAS microvessels compared to 0% KRAS microvessels (45.1±22.7 and 49.6±22.2 respectively, Figure 3B), as assessed by measuring the fluorescence intensity in the lumen and extravascular space and normalizing to the interstitial lumen intensity at 1 min. This value is displayed as a ratio of extravascular over intravascular intensity. The permeability coefficient was also calculated and indicated that there was a significant increase in vessel permeability in 10% and 25% KRAS vasculatures (14.3±8.1 and 22.7±7.6 x10^-2^ μm/s) compared to 0% KRAS vasculature (2.46±1.1 x10^-2^ μm/s) (Figure 3C). The observed increased vessel permeability in 10% and 25% KRAS vasculatures is consistent with poor vascular integrity in human AVMs.

**Figure 3:**
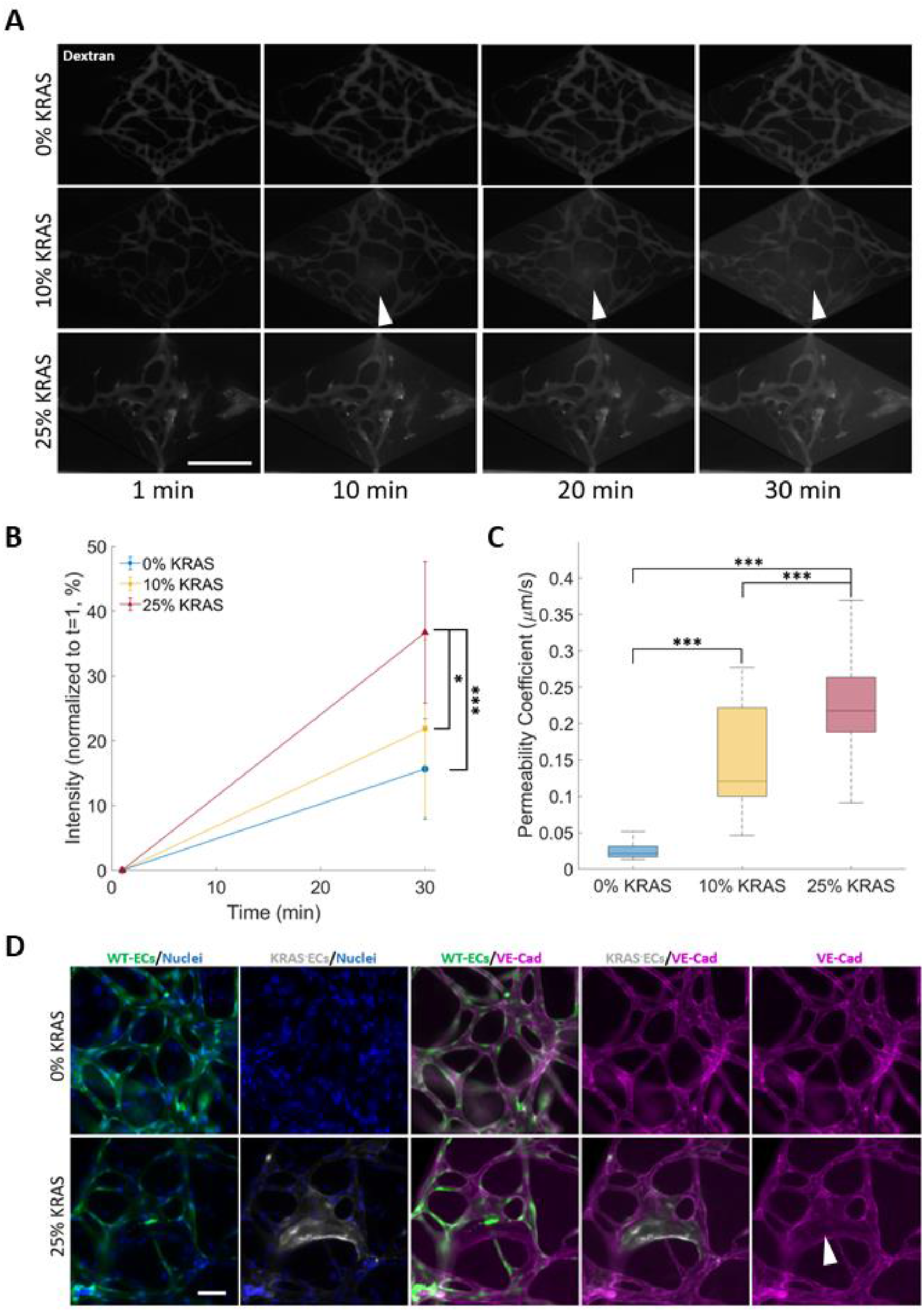
AVM-like microvessels display increased permeability. (A) Representative time course images of microvessel perfusion with 70 kDa dextran conjugated to rhodamine-B (rhod B). The 1 min timepoint denotes the moment vessels are initially fully perfused with dextran. White arrows in middle panel show leakage originating from vessel (scale bar 500 μm). (B) Quantification of microvessel 70 kDa dextran-rhod B intensity normalized to t=1 of 0% (blue), 10% (yellow), and 25% (red) KRAS4A^G12V^ microvasculatures. Fluorescence intensity was measured as a ratio of interstitial to lumen intensity then normalized to t=1. 25% KRAS4A^G12V^ microvasculatures showed a significant increase in interstitial intensity after 30 min. (C) Permeability coefficient of 0% (blue) 10% (yellow) and 25% (red) KRAS4A^G12V^ microvessel conditions. There was a significant increase in permeability as the percentage of KRAS4A^G12V^ increased. (One-way ANOVA with Tukey’s multiple comparison test; N=3, *p < 0.05, ***p < 0.001; mean ± SD) (D) Immunofluorescence images of 0% and 25% KRAS4A^G12V^ EC microvasculatures on day 7. VE Cadherin (purple) staining is reduced in areas (indicated by arrow) with KRAS4A^G12V^ EC (white) but remains intact throughout the rest of the vessel network with normal ECs (green). (scale bar 100 μm).

To investigate a potential cause of barrier dysfunction, we stained for the cell-cell adhesion molecule VE-cadherin, previously implicated in KRAS4A^G12V^ EC permeability defects in monolayers ^7,9^. Indeed, immunofluorescence staining showed that there was a reduction in VE-cadherin staining at the cell junctions of KRAS4A^G12V^ ECs (Figure 3D, arrows). The junctions of surrounding WT-ECs remained intact and unaffected. This confirmed that the increased permeability of the KRAS microvessel network is due to the disruption of adherens junctions in the KRAS4A^G12V^ ECs, thus, recapitulating the compromised barrier function associated with AVMs.

### 3.4 Improvement of vessel barrier function by treatment with a MEK Inhibitor

An important goal of a human model of AVM *in vitro* is to assess the effects of potential drugs in both structural and functional hallmarks of AVMs in a human-relevant model. We treated established 25% KRAS vasculatures with a MEK inhibitor (MEKi, SL327 ^7,9^) and assessed vessel permeability. DMSO (vehicle) and a PI3K inhibitor (PI3Ki, LY294002) acted as negative controls. PI3K is a downstream effector of KRAS in other cell types, but is not activated in KRAS4A^G12V^ ECs ^7^. To understand if MEKi would reverse established AVM defects, drug treatments started on day 5 (Figure 4A) when vessel networks were stable. Vessel permeability to 70 KDa MW dextran-rhod B was assessed on day 7 post-seeding.

**Figure 4:**
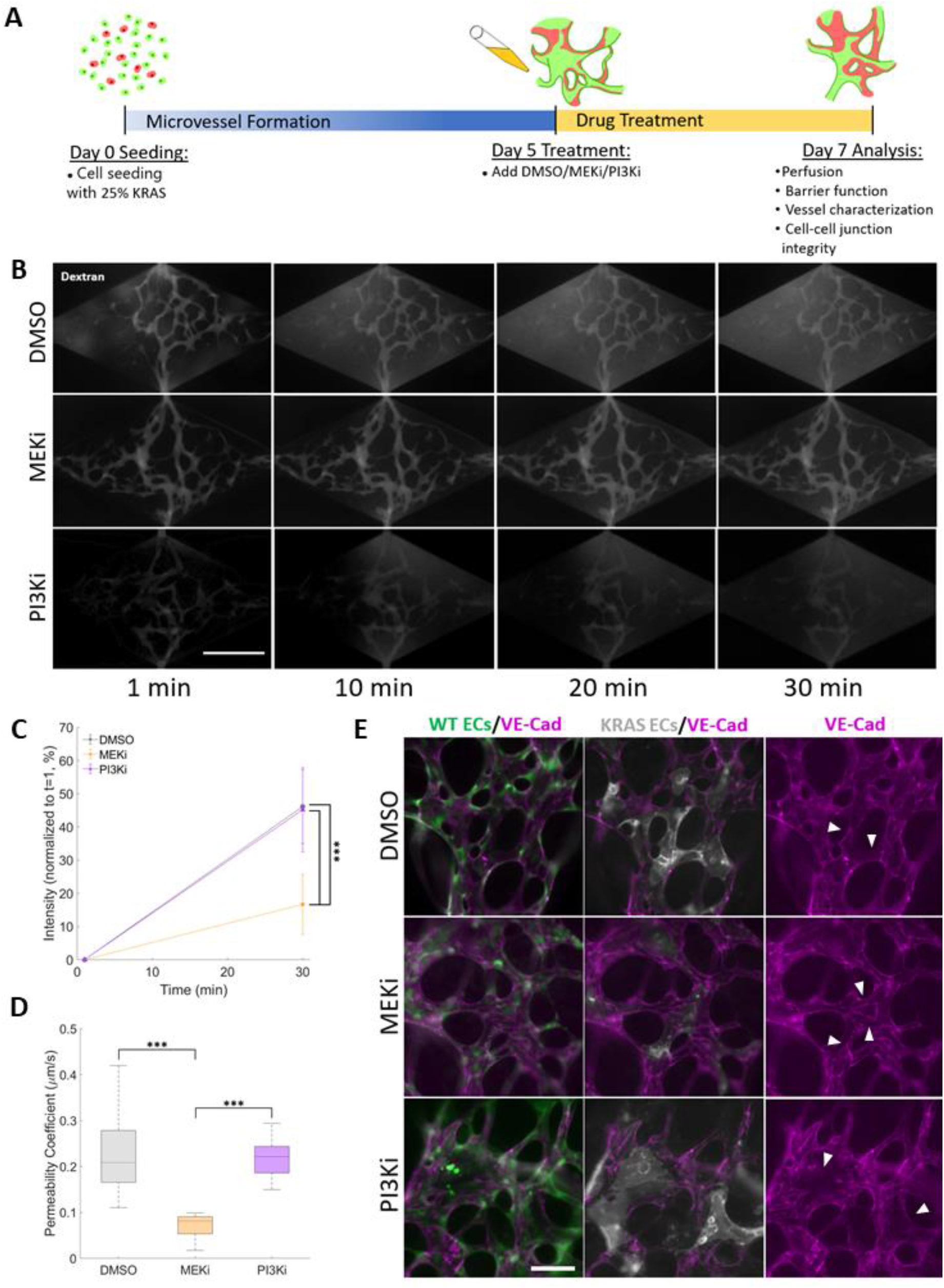
MEK inhibition restores vessel integrity of AVM-on-a-chip. (A) Schematic of drug treatment protocol. (B) Time course images of microvessel perfusion with 70 kDa dextran-rhod B (scale bar 500 μm). (C) Quantification of microvessel 70 kDa dextran-rhod B intensity normalized to t=1 of 25% KRAS4A^G12V^ microvasculatures treated with different drugs starting at day 5. Fluorescent intensity was measured as a ratio of interstitial to lumen intensity normalized to t=1. MEKi-treated AVM-like microvasculatures displayed significantly lower dextran extravasation than DMSO or PI3Ki. (D) Permeability coefficient of 25% KRAS4A^G12V^ microvasculatures treated with DMSO (gray), MEKi (orange) or PI3Ki (purple). The permeability of MEKi treatment microvasculatures was significantly lower than that of DMSO or PI3Ki treated AVM-like vessels. (One-way ANOVA with Tukey’s multiple comparison test; N=3, ***p < 0.001; mean ± SD) (E) Immunofluorescence images of day 5 treated AVM-on-a-chip. MEKi-treated AVM-like microvasculatures have an increase in VE-cadherin (purple) rich adherens junction staining in KRAS4A^G12V^ ECs (white, arrows indicate areas of increased VE-cadherin) compared to DMSO or PI3Ki treatment (arrows indicate areas of VE-cadherin disconnect) (scale bar, 100 μm).

Image analysis revealed that treatment with MEKi led to a robust reduction in interstitial fluorescence intensity of the 25% KRAS vasculatures (Figure 4B, middle panel) compared to DMSO- or PI3Ki-treated vasculatures (Figure 4B, top and bottom panels, respectively). In the MEKi-treated microvessels, there was no apparent leakage. Assessment of the normalized fluorescent intensity ratio showed a significantly lower permeability in MEKi-treated vasculature (Figure 4C, 12.5±5.8) compared the DMSO and PI3Ki treated vasculatures (Figure 4C, 44.3±12.7 and 55.2±7.5, respectively). As expected, assessment of the permeability coefficient corroborates findings from the analysis of fluorescent intensity, with the MEKi-treated networks displaying significantly lower permeability (9.0±7.0 x10^-2^ μm/s) than DMSO- or PI3Ki-treated networks (23.1±9.5 x10^-2^ and 21.6±6.4 x10^-2^ μm/s, respectively) (Figure 4D). Immunofluorescent staining of MEKi-treated vasculatures showed partial restoration of cell-cell junction localization of VE-cadherin in KRAS4A^G12V^ ECs, suggesting a role for cell-cell VE-cadherin binding in the reduction of vessel permeability. In the DMSO and PI3K treated microvessels, KRAS4A^G12V^ ECs still lacked VE-cadherin, while WT-ECs cell junctions remained intact (Figure 4E).

Next, we questioned if the restoration of barrier in AVM-like vasculatures by the MEKi would be accompanied by a concomitant decrease in vessel width. Vessel width was highly heterogeneous and there were no changes in width in response to drug treatment (Figure 5A). The average vessel width was 45.1±22.7, 46.8±18.5, 49.6±22.2 μm for the DMSO, MEKi, and PI3Ki treatments, respectively (Figure 5B), which indicates that treatment with MEKi for 2 days is not effective in reversing vessel enlargement.

**Figure 5:**
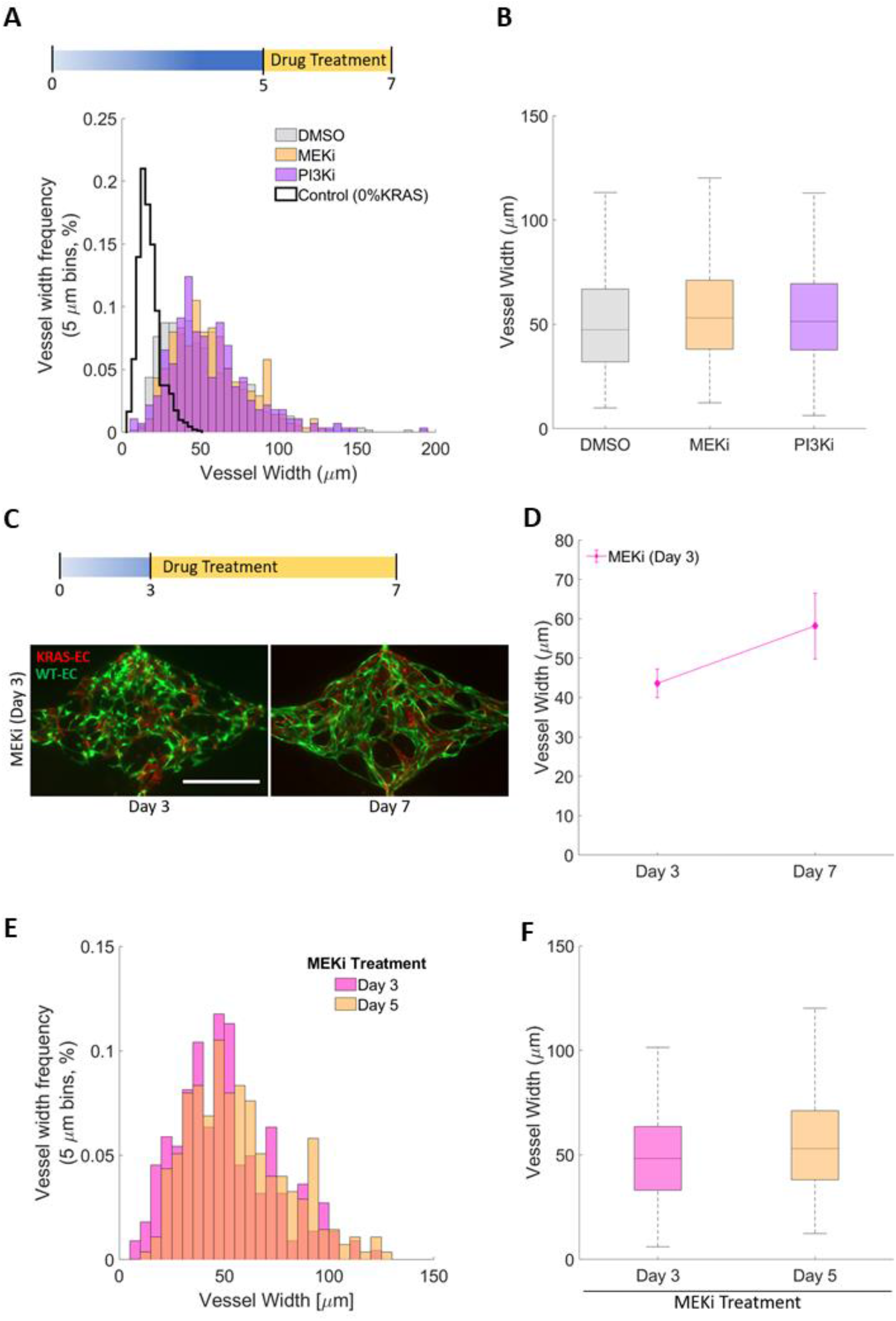
MEK inhibition does not affect width of AVM-on-a-chip. (A) Day 7 distribution of 25% KRAS4A^G12V^ microvessel widths treated with different drugs starting at day 5. (bin size: 10 μm). (B) Average vessel width at day 7 of 25% KRAS4A^G12V^ microvessel treated with DMSO, MEKi, or PI3Ki starting at day 5 show no significant differences. (One-way ANOVA with Tukey’s multiple comparison test; N=3; mean ± SD). (C) Day 7 images of KRAS4A^G12V^ microvessels treated with MEKi starting at day 3 (scale bar 500 μm). (D) Average vessel width of 25% KRAS4A^G12V^ microvasculatures before (day 3) and after (day 7) treatment with MEKi starting day 3. (Paired Student’s t-test; N=3; mean ± SD). (E) Day 7 vessel width distribution of 25% KRAS4A^G12V^ microvasculaturs treated with MEKi starting on day 3 or day 5. (bin size: 10 μm). (F) Average day 7 vessel width of 25% KRAS4A^G12V^ microvasculatures treated with MEKi starting on day 3 or day 5. (Unpaired Student’s t-test; N=3; mean ± SD).

We then tested if treatment with MEKi at an earlier time point, when the vasculature was not yet fully formed, would prevent KRAS4A^G12V^ EC-induced vessel enlargement. Therefore, we started drug treatment at day 3 post-seeding and assessed vessel width at day 7. Interestingly, treatment with the MEKi starting at day 3 did not prevent vessel enlargement (Figure 5C, D) indicating that EC assembly into microvessels was not affected by MEKi treatment (Figure 5D). In addition, a comparison of vessel widths between 25% KRAS vasculatures treated with MEKi beginning on day 3 or 5 showed that treatment with MEKi did not affect vessel width distribution (Figure 5E) and that the average vessel width at the endpoint (Day 7) was the same (Figure 5F).

## Discussion

The events that take place during sporadic AVM initiation and progression remain poorly understood. Accordingly, this lack of knowledge has stymied the development of pharmacological treatment options. In this study, we combined WT ECs with ECs harbouring a KRAS4A^G12V^ mutation and generated the first *in vitro* microfluidic model of AVMs. This heterogeneous mixture of mutant and normal ECs was chosen to reflect the heterogeneity found in clinical samples of AVMs ^7^. This platform focuses on the specific KRAS4A^G12V^ contributions to the early steps in vessel formation. The resulting microvessel network recapitulates the key hallmarks of AVM: vessel dilation, tortuosity and increased permeability. Previous studies of KRAS4A^G12V^ EC cultures reported changes in EC phenotypes, including increased cell size and irregular cell shape ^7^. Within our 3D vessel model, KRAS4A^G12V^ ECs displayed increased cell size compared to vessels formed by WT-ECs in the same microvessel network. In addition, we demonstrated that KRAS4A^G12V^-driven vessel enlargement was cell-autonomous, as mutant cell-containing vessels did not affect surrounding wildtype vessels in the same network. Additionally, defective VE-cadherin localization was observed only in KRAS4A^G12V^ ECs and not in WT-ECs. Thus, KRAS4A^G12V^ mutant ECs are necessary and sufficient for eliciting the disease phenotype. The resulting enlarged vessels also increased vessel area coverage, with significantly shorter vessel branches and an increased ratio of branches to junctions. Additionally, KRAS4A^G12V^ ECs localized preferentially to vessel junctions, resulting in vessel junctions that were much more prominent.

Previously, KRAS4A^G12V^ ECs have also been noted in monolayers^7^ for their abnormally large shape. Here we speculate that the same KRAS4A^G12V^-dependent alteration in ECs morphology is responsible for this vessel dilation. Thus, our humanized AVM model indicates that mutant KRAS enlarged vessel lumens are a consequence of increased EC size, rather than enhanced proliferation. Further investigation will determine if these enlarged KRAS-mutant vessel lumens result in increased shear stress leading to abnormal cellular mechanosensory responses or create turbulent shear to disrupt vascular homeostasis ^42^.

One limitation to note is that when measuring the branch length in the KRAS4A^G12V^ conditions, all vessels in the network were included. Some vessels may not have been KRAS4A^G12V^ positive. In addition, most vessels were likely not parallel to the XY plane, so the measured length only captures features in the XY direction. As a result, this would influence the vessel width and length measurements. Given the 100 μm height of the chamber in relation to the average vessel width that ranged from 20-60 μm, we assumed that the contribution of the Z component length would be very small and negligible.

Another key feature of the AVM-on-a-chip model is that it recapitulated increased permeability at the site of KRAS4A^G12V^ positive vessel segments due to the loss of VE-cadherin^+^ adherens junctions. Like diameter measurements, the KRAS4A^G12V^ ECs do not disrupt surrounding WT-EC cell junctions. Previous transwell leakage assays showed increase permeability due to breakdown of VE-cadherin junctions^9^. Here, we show that increased permeability is also a feature of AVM-like microvessels generated from KRAS4A^G12V^ ECs, despite the presence of luminal flow, which is a known cue for vessel barrier tightening ^43,44^.

These findings are also consistent with findings in a KRAS4A^G12V^ zebrafish AVM model^9^. However, *Nikolev et al*. demonstrated that KRAS mutant ECs comprise only a small fraction of the total ECs within AVM tissue samples^7^. A small shunt may take several years to progress into the characteristic, tangled nidus with dilated draining veins characteristic of mature, symptomatic brain AVM. Thus, it is likely that the AVM-like vasculatures formed here capture only nascent AVMs that have yet to remodel into the complex nidus that is a mosaic of wild-type and KRAS mutant ECs. An alternative explanation could be that KRAS mutations are necessary but not sufficient to induce a bAVM, and a “second hit” (e.g. from an environmental factor, or another mutation) is necessary to induce more severe presentations of the disease that is typically diagnosed in patients, as occurs in other cranial AVMs and in CCM ^45,46^. In the future, we can use our platform to investigate potential effects of increased flow rates/shear stress in AVM-like vessels. It is well known that disturbed flow, typically developed in junctions and curvature, results in EC dysfunction in vascular disease ^47^. Considering the enlarged junctions, where KRAS4A^G12V^ HUVECs preferentially reside, and irregularly shaped branches, these areas would be susceptible to creating disturbed flow and potentially a maladaptive mechanosensory response.

Some studies have suggested that AVM formation is an inappropriate angiogenic response characterized by defective vessel maturation ^10,14,48,49^. Indeed, brain AVMs have been previously shown to harbor increased expression of angiogenic genes and downstream VEGF signalling ^50–52^. KRAS^G12V^ ECs were also found to have elevated levels of VEGF and a transcriptional signature that resembles VEGF stimulated cells^7^. VEGF (originally termed vascular permeability factor, VPF) increases vessel permeability ^50^, as well as angiogenesis ^51^. This may explain the breakdown of VE-cadherin junctions that we have observed in the AVM-like microvessels, typically associated with early angiogenic stages ^55–57^. Although no overt signs of angiogenesis were seen in the AVM-like vessels (e.g. sprouting), the increase in branch number is suggestive of increased vessel formation in KRAS4A^G12V^ vasculatures.

In previous monoculture and animal models, MEKi has been investigated as a possible treatment strategy ^7,9,25,58,59^. In monocultures, MEKi redirected VE-cadherin to cell junctions and restored normal F-actin organization and cell shape by suppressing ERK phosphorylation ^7^. Transwell permeability assays with KRAS4A^G12V^ HUVECs showed reduced leakage after 18 hours of MEKi treatment. In zebrafish models, MEKi reversed vessel enlargement and hemorrhage ^9^. Mouse models of brain AVMs resulting from delivery of brain trophic AAV-KRAS^G12V^ revealed that treatment before AVM development led to significantly less brain AVMs and improved cognitive function ^25^. We have demonstrated that treatment with MEKi recovered VE-cadherin localization in KRAS4A^G12V^ HUVECs, which resulted in improved vascular wall integrity. However, treatment with MEKi did not reverse (i.e., treatment at day 5 post-seeding when microvessels are mostly formed and there is perfusion) or prevent (i.e., treatment at day 3 post-seeding prior to the formation of an interconnected vessel network) vessel enlargement. With this platform, for the first time, we were able to uncouple vessel enlargement from vascular permeability and showed that MEKi can improve permeability without affecting vessel size.

Differences in AVM formation on chip compared to the clinic, where non-mutant ECs are found to contribute to AVMs, suggest that other signals may be required ^45^. For instance, the inability of a MEKi to reduce vessel size in AVM-like vessels suggests that other cell types or signaling pathways may be affected by KRAS activation in ECs. KRAS is one of the most frequently mutated oncogenes in cancer and plays a central role in several growth factor receptor tyrosine kinase signalling pathways ^7,8,60,61^. As a result, targeting the MAPK/ERK and KRAS protein activity for cancer treatment is of high interest. Two recent studies report the off-label use of Trametinib, and FDA-approved MEKi, improved AVMs, including an overall reduction in patient AVM size ^62,63^. Using this platform, we can systematically assess and isolate external factors such as perivascular cell, inflammation, increased fluid flow, and tissue-specific interactions to determine how these parameters influence AVM initiation and progression. We speculate that reduced pericyte coverage and paracrine signalling may also play a role in decreased AVM vessel stability ^64^. In 3D collagen co-cultures of KRAS4A^G12V^ HUVECs and pericytes, the KRAS4A^G12V^ HUVECs formed abnormally wide tubes with a lack of pericyte recruitment compared to normal HUVECs. Additionally, there was a decrease of basement membrane protein deposition, suggesting a lack of pericyte-induced maturation ^65^. In the future, pericytes can be included in the microfluidic devices to better mimic vessel composition ^32,33,66^. In doing so, we can gain new insight into KRAS4A^G12V^ EC-pericyte crosstalk and whether the lack of pericyte association exacerbates AVM phenotypes ^16^.

In conclusion, we have developed and validated an AVM-on-a-chip platform using a mosaic composition of WT and KRAS4A^G12V^ ECs that is sufficient to drive AVM defects: vessel distension and hyperpermeability. These findings confirm previous results in 2D monolayer culture and animal models and provide new insight into the 3D mechanism of the KRAS4A^G12V^ mutation ^7,9^. Additionally, this platform represents an ideal method to create other vascular disease models *in vitro* using mutant EC cell lines. Several vascular malformations (venous, capillary, lymphatic, cavernous, telangiectasis, and arteriovenous) have been associated with genetic mutations ^1,67^. A hereditary form of AVMs, HHT has been linked to mutations in *BMP9/GDF2*, *SMAD4*, *ALK1*, and *ENG* ^3,7,42,43^. Additionally, there have been other mutations identified in the MAPK pathway (*HRAS*, *BRAF*, and *MAP2K1*) associated with AVM ^6,70^. Cerebral cavernous malformations (CCMs) are caused by several different mutations in *KRIT1*, *CCM1*, *23*, or the gene *PDCD10* ^46^. It is currently unclear how mutations in these disparate genes alter EC phenotype to drive vascular malformations. This platform would be of interest to gain a better understanding of the disease phenotypes resulting from these EC mutations. In this way, gene-specific differences in AVM formation and other vascular malformations may be revealed. Importantly, the platform described here can be used to test pharmacologic treatments for ameliorating these vascular malformations.

## Acknowledgements

This work was supported by grants from the Canadian Institutes of Health Research (CIHR) (CIHR PJT153160) and a Discovery grant from the Natural Sciences and Engineering Research Council (NSERC RGPIN 06621-2017) to S.S.N and grants from CIHR (CIHR PJT155922) and the U.S. Department of Defense (W81XWH-18-1-0351) to J.D.W. and J.E.F. K.S. was partially supported from the Mount Sinai Hospital Graduate Scholarship in Science and Technology. M.L. was partially supported from the NSERC CREATE Training program in organ-on-a-chip engineering and entrepreneurship (TOeP) and Wildcat Graduate Scholarship. R.W. was supported by a Canada Graduate Scholarship from CIHR. We thank Dr. R. Kamm for providing the IM-HUVECs (GFP WT-ECs) and Dr. Andreas Hayer for the hTERT-HUVECs, O. Mourad for the support with statistical analysis. K.S. conducted majority of the experiments and wrote the manuscript. M.L. contributed to image analysis and experimental repeats. R.W. developed and validated the KRAS^G12V^ inducible IM-HUVEC line and designed the MEKi experiment. W.D.T. created and validated the inducible KRAS constructs, while J.D.W. supervised W.D.T. and designed the KRAS^G12V^ inducible constructs and also revised the manuscript. J.E.F. provided supervision to R.W., designed experiments and revised the manuscript. S.S.N. provided supervision and guidance for this project, supervision to K.S. and M.L., manuscript writing and revisions.

## Manuscript Supplemental Figures Legends

**Supplemental Figure 1.**
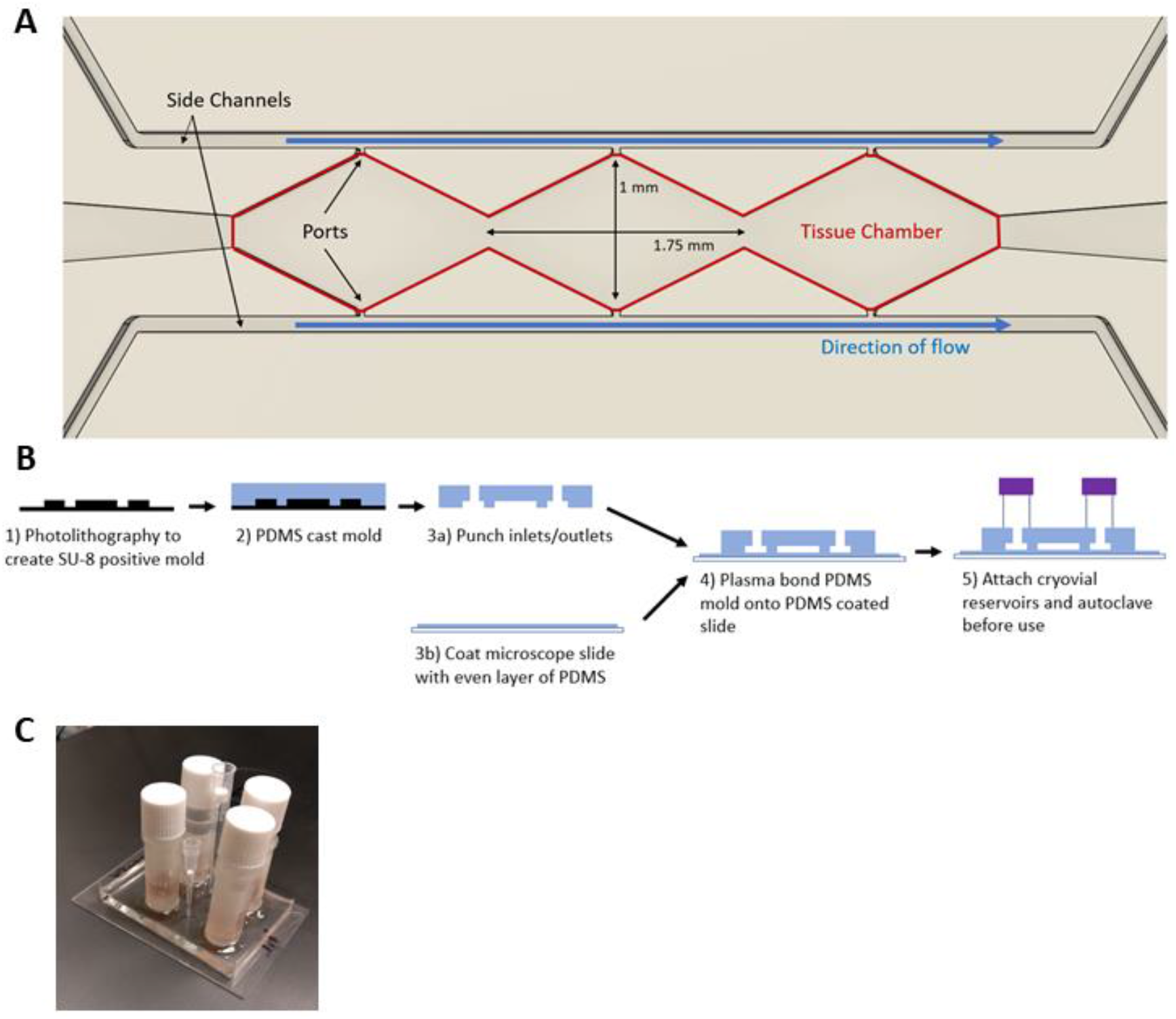
Fabrication of Devices. (A) Design of microfluidic device. Three overlapping diamond channels (tissue chamber, red) are connected to two parallel 100 μm wide side channels through a 50 μm long port. Side channels are used to perfuse media into the tissue chamber during microvessel formation. The length of media channels and gel loading channel were arbitrarily chosen for ease of handling. (B) Schematic of fabrication process. PDMS negative molds were made from silicon wafers with design etched on. 2 μm inlets were punched for the media ports and 1 μm used for gel loading. The PDMS mold was plasma bonded onto glass slide coated with PDMS. Cryovials were glued on using PDMS, acting as media reservoirs. (C) Photo of assembled microfluidic device.

**Supplemental Figure 2.**
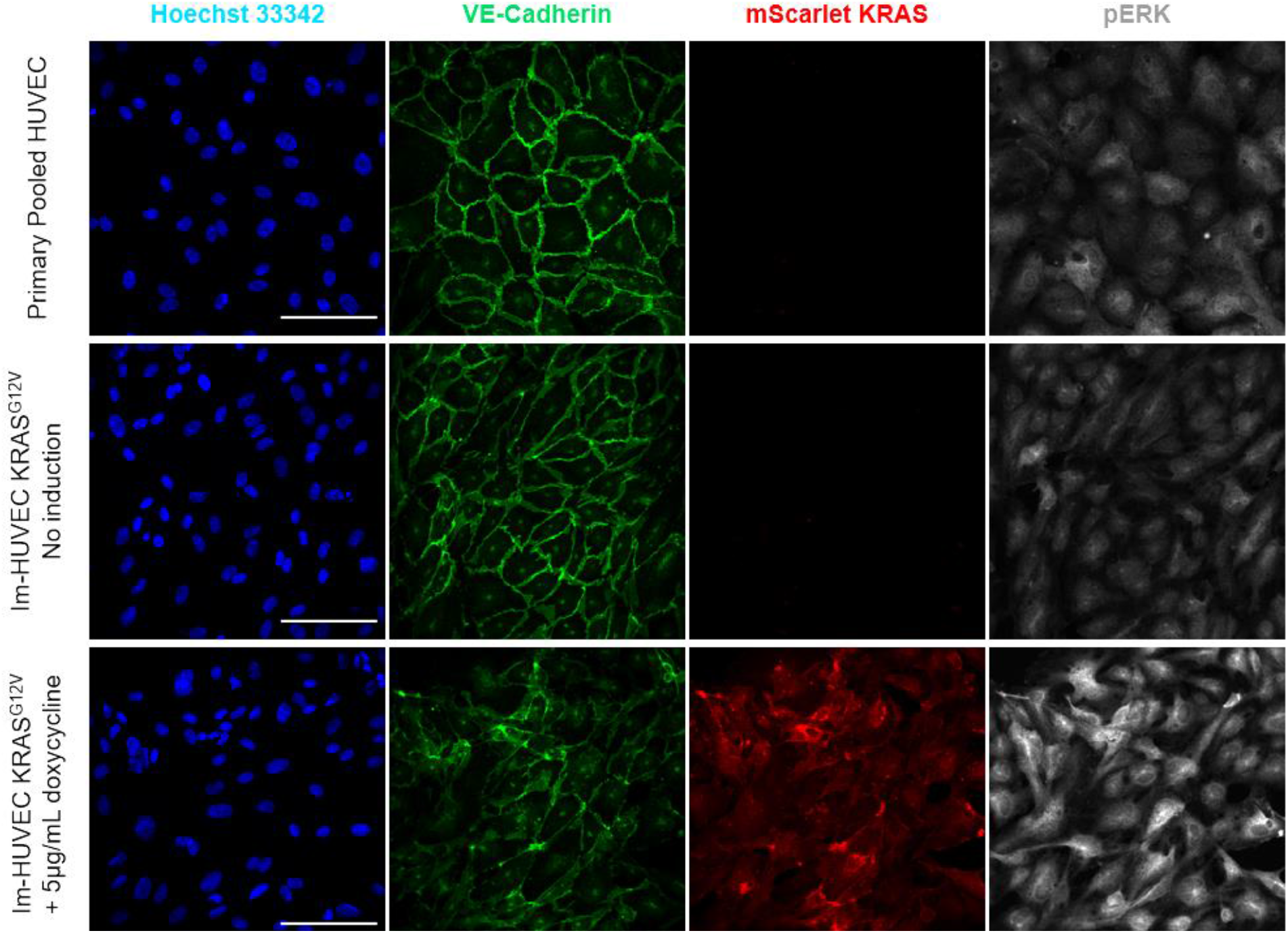
Validation of the Im-HUVEC KRAS^G12V^ cell line. Representative immunofluorescence staining of primary HUVEC and Im-HUVEC KRAS^G12V^ cells (with and without doxycycline induction) with VE-cadherin (green channel) and pERK1/2 (Thr202/Tyr204) (deep red channel). mScarlet-tagged KRAS4A^G12V^ (red channel) is under the regulation of a Tet-ON promoter and is only expressed upon doxycycline induction (5μg/mL). The images were taken at 40X magnification (scale bars, 100μm).

**Supplemental Figure 3.**
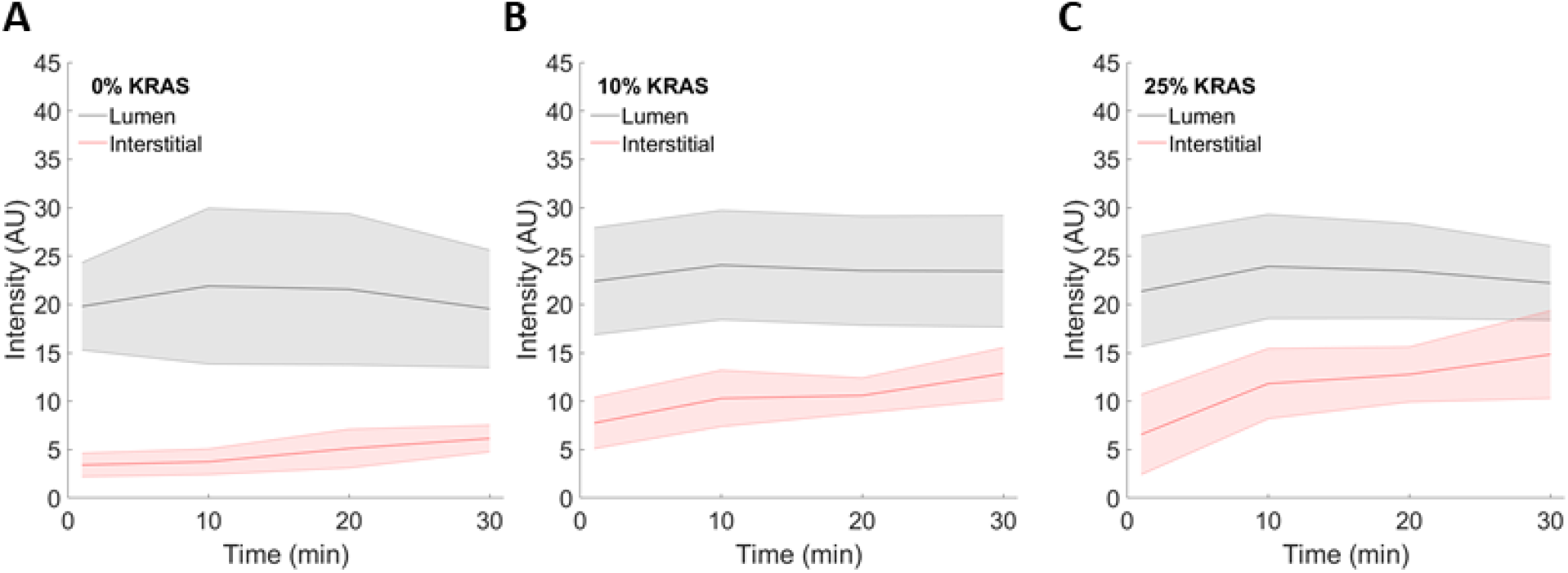
Luminal and interstitial intensity of AVM-like microvessels. Raw intensity values of vessel lumen (black) and interstitial space (red) measured at 1, 10, 20 and 30 minutes following injection of 70 kDa rhod B dextran of (A) 0%, (B) 10%, and (C) 25% KRAS4A^G12V^ microvasculatures. (N=3)

## Data Availability Statement

The data supporting the findings of this study are available from the corresponding author upon request.

